# Does urbanization lead to parallel demographic shifts across the world in a cosmopolitan plant?

**DOI:** 10.1101/2023.08.14.552623

**Authors:** Aude E. Caizergues, James S. Santangelo, Rob W. Ness, Fabio Angeoletto, Daniel Anstett, Julia Anstett, Fernanda Baena-Diaz, Elizabeth J. Carlen, Jaime A. Chaves, Mattheau S. Comerford, Karen Dyson, Mohsen Falahati-Anbaran, Mark D.E. Fellowes, Kathryn A. Hodgins, Glen R. Hood, Carlos Iñiguez-Armijos, Nicholas J. Kooyers, Adrián Lázaro-Lobo, Angela T. Moles, Jason Munshi-South, Juraj Paule, Ilga M. Porth, Luis Y. Santiago-Rosario, Kaitlin Stack Whitney, Ayko J.M. Tack, Marc T.J. Johnson

**Author notes:** Corresponding author: Aude E. Caizergues. Members of the lead team (Aude E. Caizergues, James S. Santangelo, Rob W. Ness, Marc T.J. Johnson).

## Abstract

Urbanization is occurring globally, leading to dramatic environmental changes that are altering the ecology and evolution of species. In particular, the expansion of human infrastructure and the loss and fragmentation of natural habitats in cities is predicted to increase genetic drift and reduce gene flow by reducing the size and connectivity of populations. Alternatively, the “urban facilitation model” suggests that some species will have greater gene flow into and within cities leading to higher diversity and lower differentiation in urban populations. These alternative hypotheses have not been contrasted across multiple cities. Here, we used the genomic data from the Global Urban Evolution project (GLUE), to study the effects of urbanization on non-adaptive evolutionary processes of white clover (*Trifolium repens*) at a global scale. We found that white clover populations presented high genetic diversity and no evidence of a reduction in *N_e_* linked to urbanization. On the contrary, we found that urban populations were less likely to experience a recent decrease in effective population size than rural ones. In addition, we found little genetic structure among populations both globally and between urban and rural populations, which showed extensive gene flow between habitats. Interestingly, white clover displayed overall higher gene flow within urban areas than within rural habitats. Our study provides one of the largest comprehensive tests of demographic effects of urbanization and our results contrast the common perception that heavily altered and fragmented urban environments will reduce the effective population size and genetic diversity of populations and contribute to their isolation.

## INTRODUCTION

In the past century, urbanization has rapidly modified landscapes throughout the world. Urbanization replaces natural and rural habitats with highly disturbed, human modified habitats, characterized by a high density of humans, buildings and roads, and consequently more impervious surfaces, increased pollution, and elevated habitat loss and fragmentation (Grimm et al., 2008; Liu et al., 2020). These landscape alterations influence multiple evolutionary processes, including natural selection, mutation, gene flow and genetic drift (Johnson & Munshi-South, 2017; Miles et al., 2019; Schmidt et al., 2020; Somers et al., 2004; Yauk et al., 2008). For example, we know that cities influence natural selection, yet not all species in urban habitats have adapted to these new environments (Lambert et al., 2021). The potential of a population to adapt to a new environment is heavily influenced by the availability of genetic diversity, which is in part driven by demographic changes in population size and the connections among populations. Hence, the demographic history of a population can play a major role in its past and present evolutionary trajectory, which raises concerns about the future of urban dwelling species and their conservation. How urbanization influences the evolutionary demography of populations remains poorly understood and is the focus of our study.

Habitat fragmentation and degradation can influence population demographic processes, which in turn can affect genetic drift within populations and gene flow between populations (Hanski, 1998). Fragmentation occurs when habitats are divided into patches with limited corridors for dispersal among remaining habitat fragments. For instance, when non-urban habitats are converted into buildings and roads, it splits and potentially isolates natural patches. This phenomenon can reduce the size of or even eliminate populations, and simultaneously isolate populations by limiting dispersal among them. One predicted evolutionary outcome to urban habitat fragmentation is increased genetic drift, which can have a long-term detrimental effect on a population’s fitness due to the loss of genetic diversity, elevated inbreeding, the accumulation of deleterious mutations, and reduced ability to adapt to environmental changes. The reduced gene flow that ensues from the isolation of populations is also expected to lead to increased genetic differentiation between populations (Beninde et al., 2018; Lourenço et al., 2017; Munshi-South et al., 2016). The hypothesis of increased drift and divergence of urban populations is referred to as the “urban fragmentation model”, and it is widely thought to be the most prevalent outcome of urbanization on the non-adaptive evolution of populations (Miles et al., 2019).

While the urban fragmentation model has the most support (Miles et al., 2019), urbanization can also facilitate the ecological success and evolutionary potential of some species. In particular, human commensals may thrive in urban habitats (Carlen & Munshi-South, 2021; Medina et al., 2018; Rochat et al., 2017). In such cases, urban features and human behaviour can facilitate individual movement between populations and create corridors of dispersal and gene flow, leading to higher genetic diversity within urban populations and decreased divergence between urban populations (Miles et al., 2018). Such observations have led to an alternative hypothesis – the “urban facilitation model” (Miles et al., 2019). Understanding which of these two scenarios is likely to occur for any given species is of major importance to understanding how urbanization shapes evolution, because demographic processes such as changes in population size and dispersal influence the amount of genetic variation within and between populations, and thus the evolutionary potential of populations.

Urbanization is expected to cause similar evolutionary processes and patterns across cities (Santangelo et al., 2020), since urbanization frequently leads to similar environmental changes (McKinney, 2006; Santangelo et al., 2022). While there has been a focus on studying whether urbanization causes parallel adaptive evolution to different cities (Caizergues et al., 2022; Salmón et al., 2021), there has not been the same focus on whether there could be parallel non-adaptive processes. This leads to the question: Does urbanization consistently lead to increased genetic drift within populations and divergence between populations? Some have predicted that urbanization can drive parallel non-adaptive processes (Lambert et al., 2021; Santangelo, Miles, et al., 2020), but existing tests of this prediction are inconclusive and limited in spatial scale (Beninde et al., 2018; Combs et al., 2018; Mueller et al., 2018; Theodorou et al., 2018). Hence, understanding if urbanization causes parallel evolutionary and demographic patterns among cities throughout the globe remains unresolved.

We have been using white clover (*Trifolium repens L., Fabaceae*) as a model to understand how global urbanization affects evolution as part of the GLobal Urban Evolution (GLUE) project (www.globalurbanevolution.com). Here we combine new data with the large-scale dataset generated as part of GLUE. We previously reported that white clover frequently exhibited urban-rural clines in the production of an antiherbivore defense trait, and this repeated evolution was attributed to adaptive evolution. Previous results also revealed that populations of clover consistently showed high genetic diversity and low worldwide genetic structure (Johnson et al., 2018; Santangelo et al., 2022; Wu et al., 2021), but we did not explicitly investigate how urbanization impacts genetic drift and gene flow, or the potential for parallel non-adaptive evolution across the world.

Here, we seek to understand how urbanization impacts non-adaptive evolutionary processes, including demography and gene flow, of white clover in cities throughout the world. We addressed the following questions: (1) Does urbanization repeatedly cause changes in effective population size through time thus influencing genetic diversity and inbreeding within populations? (2) Does urbanization influence gene flow, differentiation, and structure among populations? To answer these questions, we sampled and performed whole genome sequencing of over 2000 plants from urban and rural populations of white clover in 24 cities across the world, and used both population genomic analyses and demographic modeling to reconstruct urban and rural population histories. Based on white clovers’ strong positive association with human modified habitats, we predicted demographic processes and gene flow would better reflect the urban facilitation model than the urban fragmentation model.

Specifically, we predicted that *T. repens* populations thrive more in urban areas than rural areas, which would be seen as higher effective population size, higher genetic diversity, lower inbreeding, and lower population structure due to high gene flow among populations compared to rural areas.

## METHODS

### Study system and data sampling

White clover (*Trifolium repens* L., Fabaceae) is a herbaceous perennial plant native to Eurasia. It can reproduce clonally via stolons that spread horizontally on the soil surface or through sexual reproduction via outcrossing (Burdon, 1983). Each plant produces inflorescences with numerous hermaphroditic flowers that are pollinated by a diversity of bee species (Kakes, 1997). White clover is an allotetraploid (Griffiths et al., 2019) with disomic inheritance (Williams et al., 1998). Because of its ability to fix atmospheric nitrogen, white clover has been introduced to all inhabited continents in the past several hundred years as livestock fodder and as a cover crop (Kjærgaard, 2003). Its global distribution covers a wide range of climates, and the fact that it grows in anthropogenically modified habitats (e.g., mowed grass, pastures), makes it an ideal model to study urban evolutionary biology at a global scale.

Our study expands upon the GLUE project that aimed to examine how global urbanization affects parallel adaptation (Santangelo et al., 2022). As a brief background, scientists from around the world sampled 20-50 populations from each city along urban-rural transects. In total, 110,019 plants from 6,169 populations were collected in 160 cities from 27 countries. Among these, a subset of 24 cities, chosen to capture variation in geography and climate, were subject to whole genome sequencing. For each of these selected cities, ∼42 individuals from the five subpopulations closest to the city center and an equivalent number of individuals from the five furthest rural subpopulations were selected for subsequent genomic analyses (mean ± SD = 83.8 ± 14.6 individuals/city).

While the initial analyses focused on adaptive parallel evolution of a single trait (i.e. ability to produce hydrogen cyanide) and the loci controlling it (*CYP79D15* and *Li*), here we specifically examined how urbanization affects non-adaptive genomic evolution. We used an improved and larger dataset including 19 cities from the initial GLUE project for which good quality genomic data were available (removing cities where more than 50% of the sampled individuals had a coverage <0.5X) and five newly sequenced cities (Palmerston North, New Zealand ; Punta Arenas, Chile ; Sapporo, Japan ; Vancouver, Canada ; and Warsaw, Poland), for a worldwide sample across all inhabited continents (Figure 1). The final whole genome sequence dataset included 6 European, 3 Asian, 1 African, 7 North American, 4 South American and 3 Oceanian cities, for a total of 2,013 individuals.

**Figure 1:**
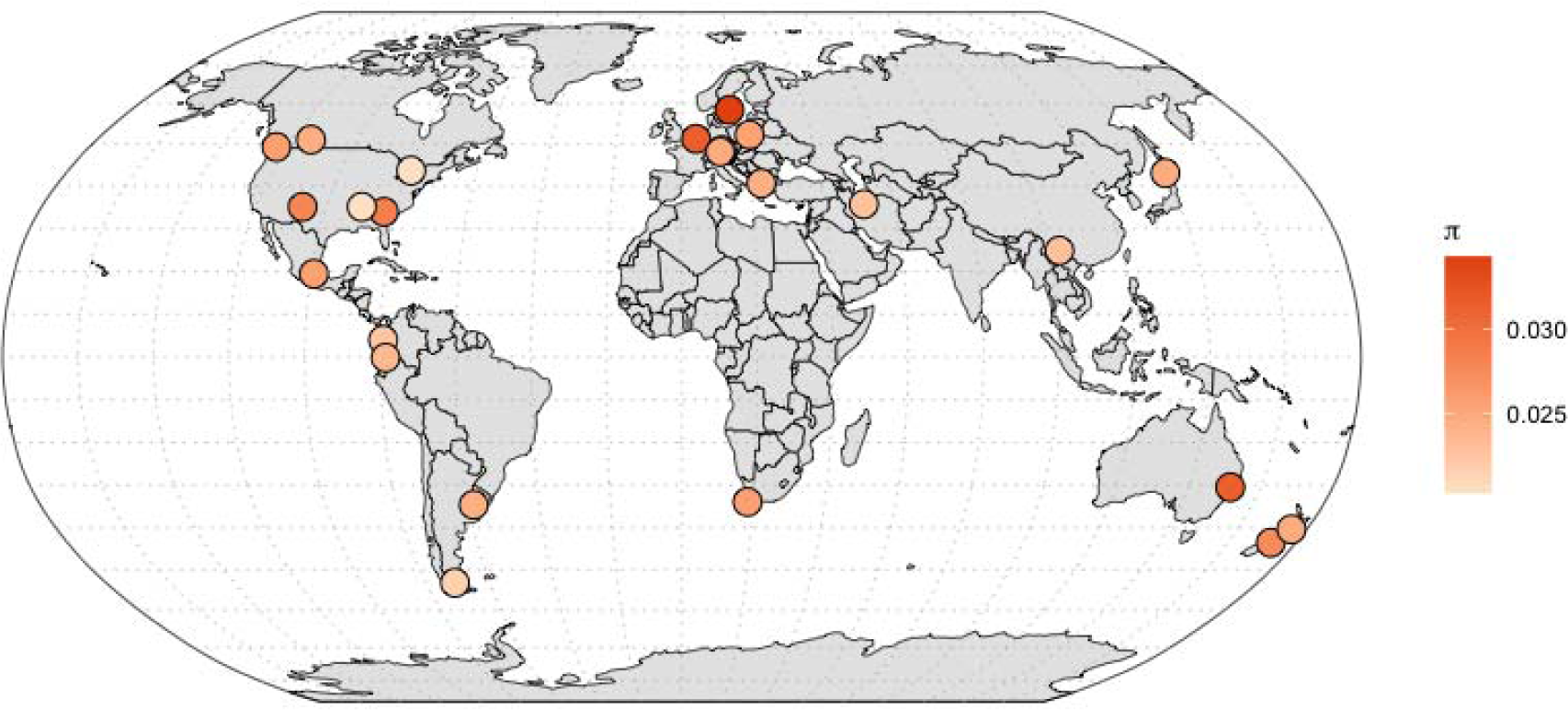
Cities sampled for rural and urban populations of white clover *Trifolium repens*.

### Molecular biology and genome sequencing

To obtain our genomic dataset, we used the sequencing protocol from GLUE (Santangelo et al., 2022) when sequencing the 5 cities newly added to the original genomic dataset. Briefly, we extracted genomic DNA from freeze-dried samples using a modified phenol-chloroform extraction protocol (detailed in Santangelo et al. 2022, Supp Text S5), and quantified DNA after extraction using the dsDNA HS Assay Kit (Fisher Scientific, Mississauga, Canada). We generated dual-indexed genomic DNA libraries following Santangelo et al. (2022) and Glenn et al. (2019), and sequenced genomic libraries with a concentration of DNA >=0.8 ng/µL. The genomes of 1,922 individuals from 23 cities (all except Toronto) were sequenced on a Novaseq 6000 S4 platform using 150 bp paired-end reads at low coverage (1.05X on average). Ninety plants from Toronto were sequenced at ∼13X as a part of another project, and downsampled using SAMtools (v1.10) to ∼2.5X to be included in this study. Seven cities from Santangelo et al. (2022) were removed from the dataset because of low sequence quality and/or low sample size (Bogota and Medellin, Colombia; Canberra and Melbourne, Australia; Hiroshima and Kyoto, Japan; Paris, France), leaving us with data from 24 cities (Figure 1).

### Sequence alignment and genotype likelihoods

Beginning with fastq sequence data, we first used *fastp* V0.20.1 to trim raw reads with the *-trim_poly_g* argument to remove polyG tails commonly generated by the Novaseq platform and performed quality checks on both raw and trimmed reads of every sample using *FastQC* v0.11.9. We then mapped trimmed reads to the *T. repens* reference genome (NCBI BioProject number PRJNA523044, Griffiths et al., 2019) with *BWA MEM* v0.7.17. We used *SAMtools* to sort, index and mark duplicates in BAM files. We performed a quality check of mapped reads with *Qualimap* v2.2.2, *Bamtools* v2.5.1, *BamUtil* v1.0.14 and *multiQC* v1.11. Since calling individual genotypes from low coverage data can lead to bias in variant detection (Han et al., 2014; Nielsen et al., 2012), we computed genotype likelihoods with *ANGSD* v0.933 (Korneliussen et al., 2014) using the *SAMtools* genotype likelihood model.

We filtered for a minimum phred-scaled base quality score of 20 and minimum mapping quality of 30. While the framework of our bioinformatic pipeline is built upon the earlier Santangelo et al. (2022) pipeline (https://github.com/James-S-Santangelo/glue_pc), all analyses and results are new, using a larger and improved dataset to specifically focus on the evolutionary signatures of population demographic change in response to urbanization. A diagram summarizing all the bioinformatic steps is available in supplementary materials Figure S1 and all analyses were integrated into a reproducible *Snakemake* (Mölder et al., 2021) pipeline (https://github.com/AudeCaizergues/glue_demography/).

The site frequency spectra (SFS) was the basis for most of the analyses performed in *ANGSD* and other software. We estimated the SFS at 4-fold degenerate sites in the *T. repens* reference genome using the “*degeneracy*” pipeline (github.com/tvkent/Degeneracy). We retained an average of 2,110,100 4-fold degenerate sites per city per habitat after filtering (see below). Since related individuals can bias population genomic analyses, and clover has the ability to grow clonally, we identified closely related individuals using *NgsRelate* (Hanghøj et al., 2019) with default parameters, and removed individuals with a pairwise relatedness r_xy_>0.5, where r_xy_=0.5 corresponds to a parent-offspring or full-sibling relationship; a total of 28 individuals were removed from subsequent analyses. After removing related individuals, we estimated the folded SFS for each urban and rural population. We also estimated the folded two-dimensional (2D) SFS for each urban-rural pair per city using the *realSFS ANGSD* function.

### Data analyses

#### Genetic diversity estimates and demographic inference

To investigate the effects of urbanization on neutral evolutionary processes within populations, our first step was to estimate genetic diversity, effective population size, gene flow and levels of relatedness within each habitat (urban and rural) of a city. To characterize population genomic parameters of diversity, we estimated nucleotide diversity based on π (pairwise nucleotide diversity), Watterson’s theta (θ_w_) (Nei, 1975; Tajima, 1983) and Tajima’s D (Tajima, 1989), from the SFS using the *thetaStat* function of *ANGSD*. Inbreeding levels within subpopulations were estimated as mean r_xy_ (i.e., the pairwise relatedness, Hedrick & Lacy, 2015), computed with *NgsRelate* using default parameters. To understand whether these diversity parameters differed between environments we included them in linear mixed models with habitat (urban vs. non-urban) as an explanatory variable and city as a random effect. We then estimated the effective population size (*N_e_*) in each habitat. Since urban populations are unlikely to be at mutation-drift equilibrium, statistics like π and θ_w_ will not capture recent changes in *N_e_*. We therefore used a model-based approach to estimate the dynamics of contemporary *N_e_*. In each city, we reconstructed variation of *N_e_* through time using *EPOS* (Lynch et al., 2020), with 1000 bootstrap iterations and a mutation rate of 1.8×10^-8^ (Griffiths et al., 2019). Raw outputs of *EPOS* were converted to a plottable format using the *epos2plot* function. These analyses gave us a detailed understanding of how urbanization affects genetic diversity and effective population size within populations.

#### Genetic structure

We next investigated how urbanization influenced population genetic structure and gene flow of *T. repens* within and between urban and rural habitats. First, since linkage disequilibrium can bias genetic structure analyses, we identified linked positions among the 4-fold degenerate SNPs with a minor allele frequency (MAF) ≥ 0.05 using *ngsLD* (Fox et al., 2019), and pruned the linked SNPs within 20kb using a r^2^ cutoff of 0.2. The pruned dataset was used for subsequent Principal Component Analysis (PCA) and admixture analyses only. To describe the genetic structure both between habitats and cities, we performed a PCA with *PCAngsd* (Meisner & Albrechtsen, 2018) that estimates a variance-covariance matrix of allele frequencies directly from genotype likelihoods. We then estimated population differentiation using Hudson’s *F*_ST_ separately for each city (Hudson et al., 1992). We computed *F*_ST_ with the *realSFS fst index* function from the 2-dimensional SFS. First, to understand genetic structure between habitats, we computed *F*_ST_ between each urban-rural pair of populations. Second, to further analyze patterns of differentiation within habitats, we computed *F*_ST_ between all pairs of subpopulations per city, and obtained an average *F*_ST_ for urban-urban, urban-rural or rural-rural comparisons separately. As a complementary analysis of urban-rural differentiation, we estimated admixture proportions for each pair of urban-rural populations with *NGSadmix* (Skotte et al., 2013). We ran the analysis for cluster numbers (K) from 2 to 10, with 10 iterations per K and selected the best K for each city using the method of Evanno et al. (2005) implemented in *CLUMPAK* (Kopelman et al., 2015). Finally, we used *GADMA* (with *dadi* engine) to estimate migration rates between urban and rural populations (Gutenkunst et al., 2010; Noskova et al., 2020). To have a more comprehensive understanding of gene flow we also estimated migration rate (*m*) using Wright’s equation *N_e_m=*(1/*F*_ST_-1)/4 (Wright, 1984).

## RESULTS

### Does urbanization repeatedly affect genetic diversity, relatedness within populations and effective population size?

Demographic processes frequently differed among cities, yet genetic diversity and effective population size were, on average, similar between urban and rural populations. While some cities displayed higher nucleotide diversity than others (e.g. see Linkoping, Sweden, Figure 2A), within cities, both urban and rural clover populations had high nucleotide diversity (mean ± SD π = 0.025 ± 0.004, Figure 2A; mean ± SD θw = 0.028±0.006, Figure S2), which did not differ between urban and rural habitats (habitat effect for π : F_1,23_=0.505, P=0.485; habitat effect for θ_w_ : F_1,23_=1.303, P=0.265). Tajima’s *D* varied among cities from -0.682 to 0.318 (Figure 2B), and was negative on average, which is consistent with recent population expansions. Tajima’s *D* also varied within cities, with 58.3% (N=14) of urban populations displaying higher *D* than their associated rural counterparts, and 41.7% (N=10) showing the opposite pattern, but there was no consistent difference between urban and rural habitats (F_1,23_=2.935, P=0.100). Such high levels of genetic diversity are consistent with large effective population sizes.

**Figure 2:**
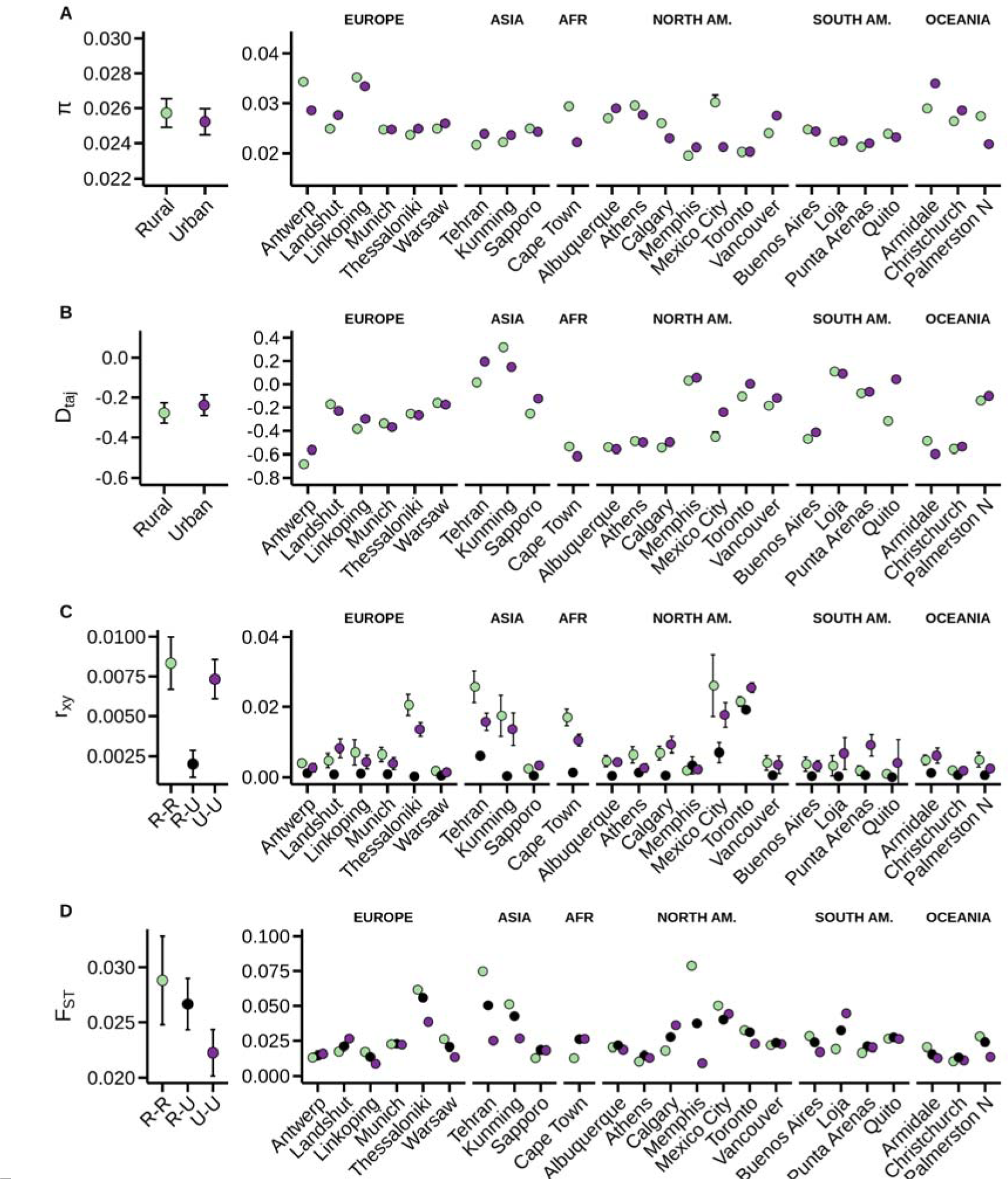
Summary of population genetic parameters averaged over all cities (left, mean±95%CI) and detailed per city (right, mean±95%CI) for: A) pairwise nucleotide diversity, B) Tajima’s D, C) averaged relatedness between individuals, and D) F_ST_ averaged between pairs of sub populations. Left graphs represent averaged values per habitat for F_ST_ and r_xy_ and genome wide averages for π and Tajima’s D. In A and B, green represents rural habitat and purple represents urban habitat. In C and D, green, black and purple respectively represent rural-rural, rural-urban and urban-urban for F_ST_ and r_xy_.

Overall, *N_e_* was high but varied substantially between cities and was more likely to decline in rural than urban habitats. *N_e_* varied among cities by four orders of magnitude, ranging from 1,750 to 40,800,000. Comparing urban and rural habitats, *N_e_* was twice as likely to be higher in the urban habitat; there were 7 occurrences of *N_e_* _urban_>*N_e_* _rural,_ (e.g. Munich, Germany, Figure 3A), 3 occurrences of *N_e_* _urban_<*N_e_* _rural_ (e.g. Toronto, Figure 3B) and 14 *N_e_* _urban_=*N_e_* _rural_ (e.g. Kunming, Figure 3C). When *N_e_* was compared between habitats with parametric (LMER) and non-parametric (Wilcoxon rank test) analyses, we found no consistent effect of urban/rural habitat on *N_e_* (LMER: X^2^ =1.181, P=0.277; Wilcoxon rank test: P=0.107). We found low levels of relatedness between individuals, and relatedness did not differ between urban and rural habitats (Figure 2C, Wilcoxon rank test: P=0.303), suggesting urbanization did not affect the propensity of inbreeding. Using the whole genome dataset, we modeled recent changes in *N_e_* through time in both habitats. Although *N_e_* did not consistently differ between habitats, urban populations were less likely to show a recent decrease in *N_e_* in the last 500 years compared to rural populations. Specifically, *N_e_* decreased in 4 urban populations, whereas it decreased in 11 rural habitats, and the remaining populations were stable between habitats (Figure S3; ^2^=4.751, P=0.029). Taken together, our results suggest that while clover shows overall high *N_e_* that maintains substantial genetic diversity within populations, urban habitats are more likely to maintain large and stable populations than rural areas.

**Figure 3:**
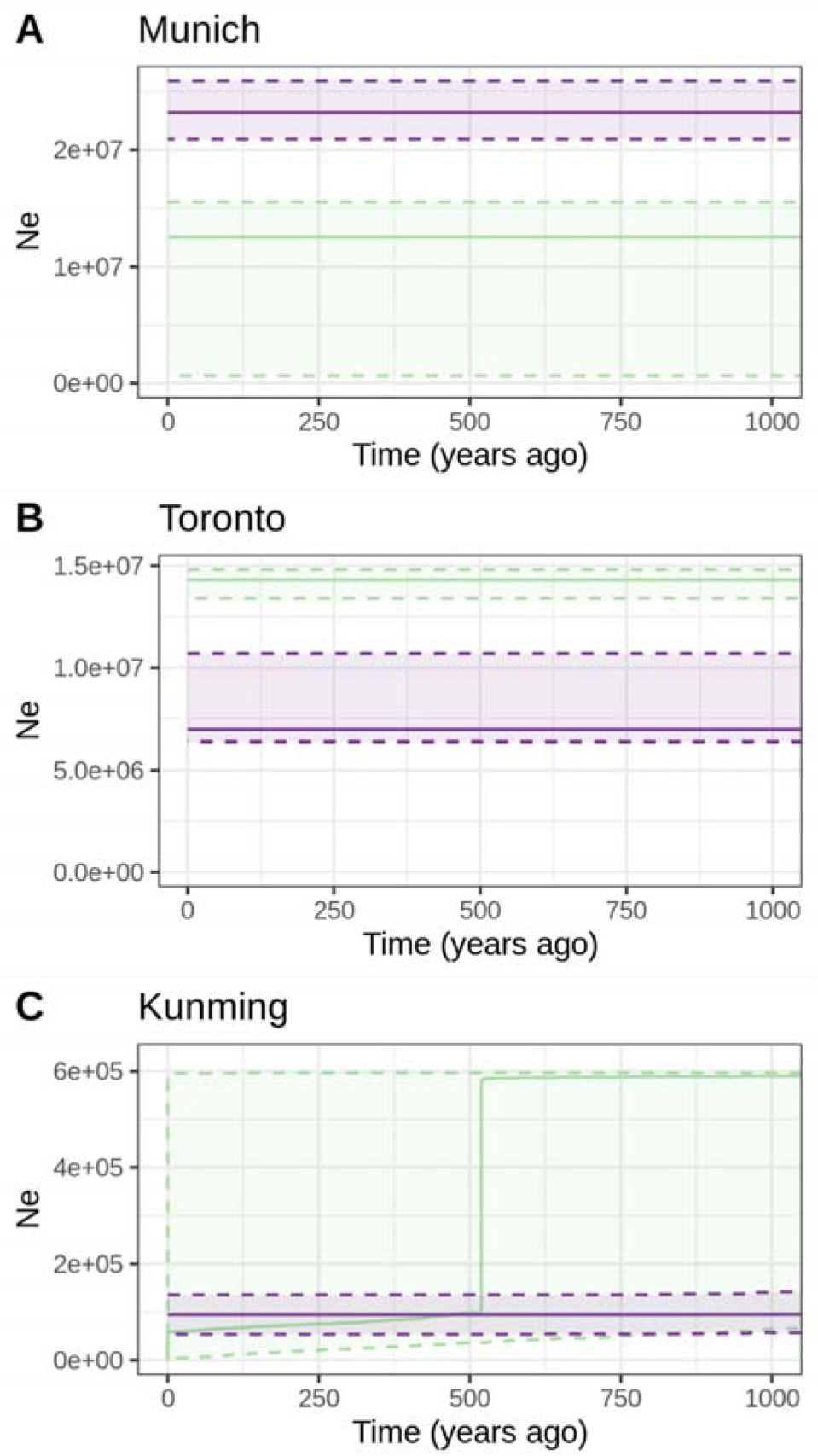
Effective population size (*N_e_*) in the past 1000 years for three representative pairs of urban-rural populations in Munich (Germany), Toronto (Canada), and Kunming (China), showing the diversity of patterns of recent *N_e_* variations across the dataset, where we find no consistent effect of urbanization on *N_e_*. Solid lines represent the median and dashed line the 5% and 95% quantiles.

### Does urbanization influence gene flow, differentiation and genetic structure?

Analysis of population structure shows that while cities can be genetically differentiated from one another on a global scale, urban and rural populations show limited structure within a given city. At a global scale, PCA analyses revealed that many cities cluster close to each other if they are geographically closer to one another. For instance, North and South American cities were typically more genetically similar to one another than they were to European cities (Figure 4 A and B). By contrast, several cities were genetically distinct, such as Cape Town, South Africa, the only African city sampled, and Tehran, Iran, and Thessaloniki, Greece, which are geographically isolated from other sampled points (Figure 4 A and B). Within a city, PCA revealed no strong pattern of differentiation between urban and rural populations (Figure S4).

**Figure 4:**
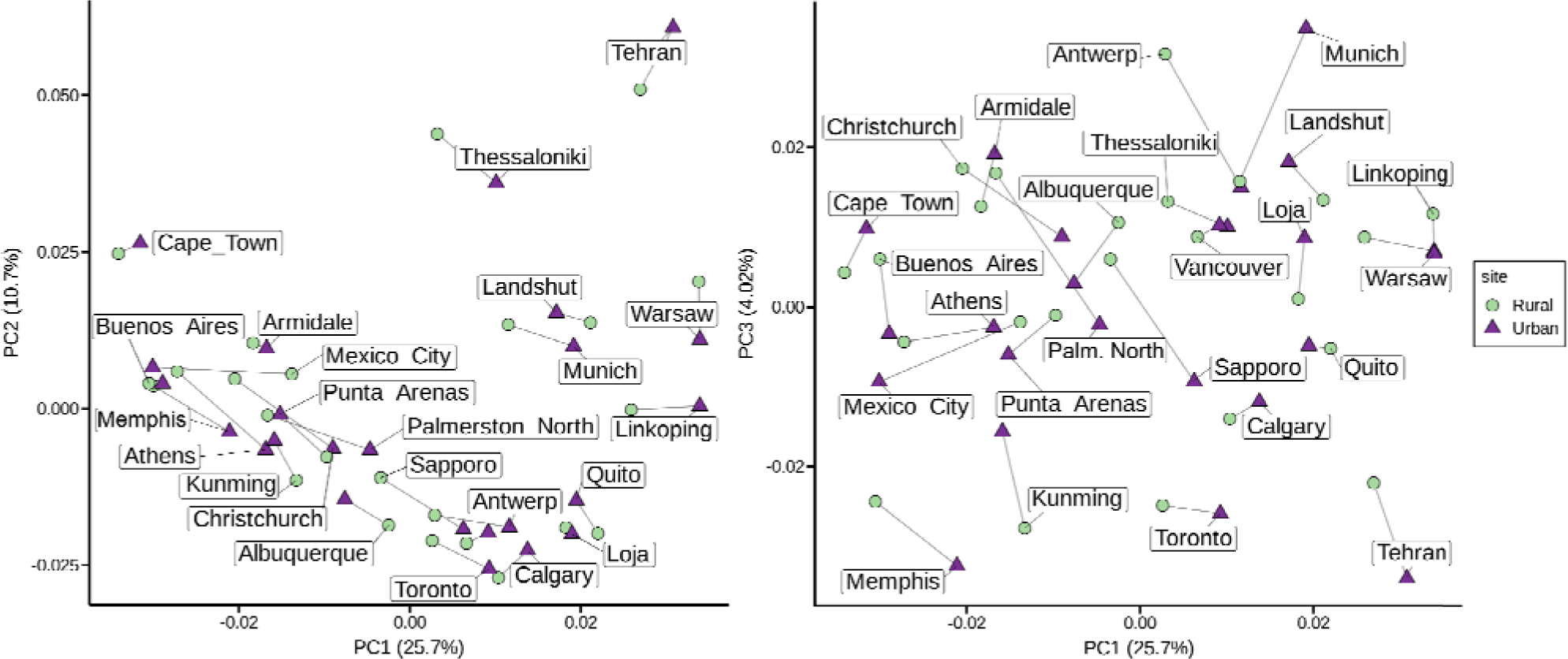
Genetic structure among *T. repens* populations revealed by a principal component analysis (PCA) on genotype likelihoods of 151,076 4-fold positions, A) on PC1 and PC2, and B) on PC1 and PC3. Each point represents the centroid of rural (green round) and urban (purple triangle) populations for each city sampled.

When population differentiation was quantified with *F*_ST_, we observed that overall between-population *F*_ST_ was low (Figure 2D), but there was evidence of fine-scale differences when moving from between-to within-habitat comparisons. Specifically, while the average *F*_ST_ between urban and rural populations was similar to *F*_ST_ within rural habitats (mean ± SD *F*_ST urb-rur_ = 0.029 ± 0.028, *F*_ST rur-rur_=0.026±0.016, Wilcoxon rank test: P= 0.931), Fst within urban habitats was 19% lower than *F*_ST_ within rural habitat, and 27% lower than than *F*_ST_ between habitats (mean±SD *F*_ST urb-urb_=0.021±0.013; Wilcoxon rank test *F*_ST urb-urb_-*F*_ST rur-rur_: P= 0.031, Wilcoxon rank test *F*_ST urb-urb_-*F*_ST urb-rur_: P= <0.001). These differences in population differentiation between habitats are consistent with more extensive gene flow among sites within urban habitats compared to the gene flow among sites within rural habitats, or between urban and rural habitats.

Estimates of historical and recent demographic processes reveal extensive gene flow between urban and rural populations. Admixture analyses identified admixture between urban and rural populations, with 2 to 6 populations (best supported “K”) found within cities that were broadly shared between urban and rural individuals, and revealed some small scale structure between populations but no strong differentiation between urban and rural habitats (Figure 5). Between habitat migration rates estimated using *dadi* showed no difference in urban-to-rural and rural-to-urban migration rates (Figure S5). Similarly, estimates of *N_e_m* (Wright 1931), showed no clear variation between habitats (Figure S6). Hence, the low genetic structure observed between habitats is likely due to high bidirectional gene flow between urban and rural habitats.

**Figure 5:**
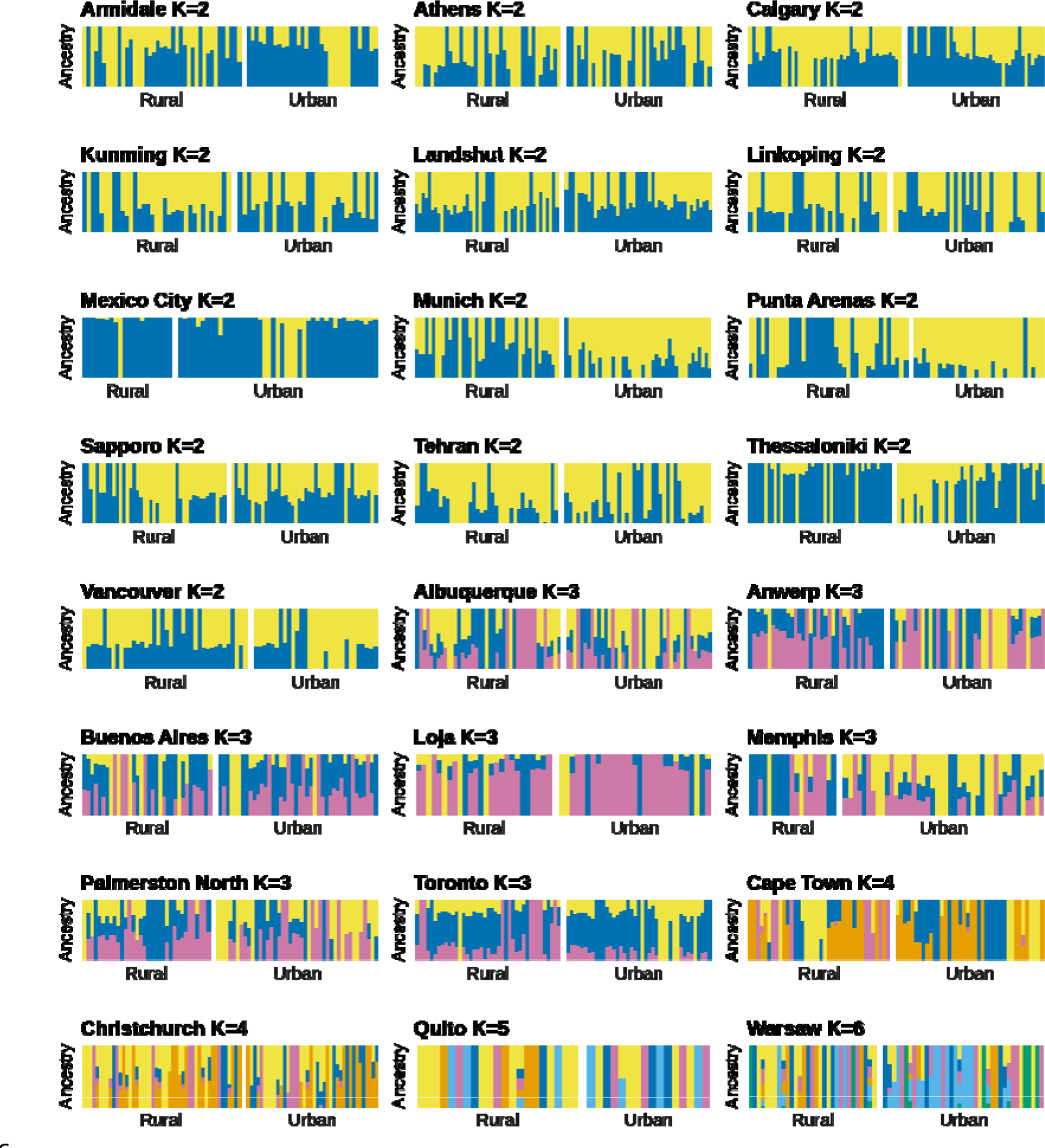
Admixture proportion between urban and rural habitat for each city estimated with *NGSadmix*. The most well supported K value for each city was selected with the method of Evanno et al. (2005). Each vertical bar represents an individual and the height of the colour indicates the proportion of an individual’s genome derived from each of the K ancestral populations within a given city.

## DISCUSSION

We tested whether global urbanization leads to parallel non-adaptive evolutionary processes and patterns in the cosmopolitan plant white clover (*T. repens*). Specifically, we sought to understand whether urban and rural habitats consistently differed in genetic diversity, inbreeding, effective population size, genetic structure and gene flow. Our genome wide analyses consisting of 1,922 individuals from 24 populations/cities revealed high levels of diversity in both urban and rural populations of white clover. Almost all urban populations had high stable *N_e_*, which was not the case for some rural populations, which disproportionately showed decreasing effective population sizes. We also detected low genetic structure among urban and rural populations and high admixture, consistent with frequent gene flow both within and among habitats. Gene flow appears to be more prevalent among urban than rural populations, leading to greater genetic homogenization in urban habitats. Our results suggest that white clover demography follows the urban facilitation model more than the urban fragmentation model, in that white clover thrives in urban environments and maintains high evolutionary potential of populations.

### Effects of urbanization on non-adaptive evolutionary processes within populations

Urbanization frequently leads to fragmentation, which often causes a decrease in effective population size and genetic diversity within populations (i.e. the Urban Fragmentation Model). This prediction has been supported by a broad diversity of studies. For example, fragmented forest habitats lead to a decrease in genetic diversity in red-backed salamanders (*Plethodon cinereus*) in the city of Montreal (Noël et al., 2007). Similarly, white footed mice (*Peromyscus leucopus*) showed reduced diversity along an urbanization gradient in New York (Munshi-South et al., 2016). Our results do not match this expected pattern of the urban fragmentation model.

Consistent with the urban facilitation model, we found that urban white clover was able to maintain high levels of genetic diversity and high effective population size throughout the sampling range of our study. Our results of high genetic diversity in urban and rural populations are consistent with previous smaller scale studies that used microsatellite markers, which showed that genetic diversity did not vary along urbanization gradients in multiple cities in Ontario, Canada (Johnson et al., 2018). These findings may also help explain how white clover has been able to rapidly adapt to repeated urbanization gradients throughout the world (Santangelo et al., 2022). Adaptive evolution is expected to be more efficient when *N_e_* is high and selection is strong. We confirmed large *N_e_* in most cities, which is consistent with the observation that white clover is frequently abundant and widespread throughout cities. A novel result of the current study is that urban habitats are more likely to maintain high and stable population sizes than rural areas. Given that urban habitats frequently undergo rapid environmental change, we may expect urban populations to experience stronger selection than rural habitats.

There is mounting evidence that the urban facilitation model, supported by our study, may apply to many species. For instance, brown rats display high genetic diversity in urban areas that can be attributed to large population sizes (Combs et al., 2018). Similarly, urban great tits (*Parus major*) showed higher genetic diversity in cities than in the associated forests (Björklund et al., 2009). White clover has many traits (e.g., clonal growth, prostrate growth habit, ability to regrow following cutting, polyploidy) that makes it well-adapted to human dominated landscapes like lawns and well-grazed pastures. In particular, clover is a crawling plant making it a poor competitor for sunlight, but it has the ability to quickly regrow following damage, and might thus benefit from mowing. Urban areas display a multitude of well mowed green spaces (parks, lawns, private backyards) where clover can grow. In addition, clover is sensitive to drought and benefits from irrigation provided to many urban spaces. By contrast, rural areas are frequently unmowed and unwatered, which can lead to clover being outcompeted and experience low fitness. Finally, urban white clover’s ability to maintain constant high population sizes compared to rural ones, might also benefit from frequent introductions of clover to cities (via seeds in sod and turfgrass) from diverse seed stocks leading to elevated diversity and low genetic structure. Hence, both the life history traits of clover and its close link with intense human activities make it a species well adapted to performing best in anthropogenic habitats.

### Effects of urbanization on genetic structure and gene flow between populations

Consistent with the urban facilitation model, we found that white clover displays high gene flow between urban and rural areas, as suggested by low levels of differentiation, limited genetic structure, high admixture proportions, and high estimates of dispersal. High levels of gene flow between populations are consistent with high genetic diversity and large population sizes, as connectivity between populations enhances allele movement and reduces genetic drift. A major element of white clover history is that it was often intentionally planted by humans as a fodder crop, to enrich soil nitrogen in rotation farming, unintentionally introduced via sod or turfgrass, or intentionally planted as clover lawns (Kjærgaard, 2003). Thus, white clover was potentially introduced multiple times in each area and moved among locations by people. Such repeated introductions have likely played a major role in the low genetic structure observed between habitats and evidence of substantial admixture.

While urban and rural clover display similar low levels of relatedness, we found the lowest differentiation among urban populations (i.e. lower within urban F_ST_ than within rural F_ST_), suggesting the urban landscape facilitates gene flow. As an obligate outcrosser, white clover depends on pollinators for its reproduction. Cities can harbor more pollinator diversity than rural areas (Wenzel et al., 2020), and since movement of pollinators is not necessarily constrained by the urban landscape (Theodorou et al., 2018), higher gene flow within cities might be driven by pollinators. For instance, in Toronto, Canada, previous research showed that pollinator visitation rate to white clover was higher in urban sites than in rural sites (Santangelo et al., 2020). This novel result of high gene flow in urban versus rural habitats, also suggests that any adaptations that do arise (e.g. HCN production, Santangelo et al., 2022) are likely to spread more quickly among urban than rural populations.

Recent literature suggests that several species may exhibit extensive gene flow in urban areas. For instance, highly mobile species such as the feral pigeons (*Columba livia*) can display high gene flow in urban areas (Carlen & Munshi-South, 2021). In small mammals, Combs *et al*. (2018) found high genetic diversity in brown rats (*Rattus novegicus*) in four cities, likely due to high gene flow and large population sizes. However, high gene flow in urban habitats is not limited to highly mobile species. In fact, species that are not necessarily good dispersers, like the western black widows (*Latrodectus hesperus*), can directly benefit from human movement and transportation networks for their dispersal (Miles et al., 2018). As a result, species that are intentionally or unintentionally moved by humans are likely to display higher levels of gene flow. Our results on white clover contribute to the growing evidence that urbanization can facilitate gene flow and reveal that such processes can be repeated across cities worldwide.

### Future directions for the study of parallel genomic evolution in cities

Cities offer a great opportunity to study how anthropogenic disturbances affect evolutionary processes at a global scale. In line with our study, there has been a recent focus on studying if similar environmental changes caused by urbanization lead to independent, parallel evolutionary trajectories. Most of the recent focus has been directed towards studying parallel adaptive phenotypic shifts. While common garden or experimental approaches can provide useful insight on the parallelism of responses to urbanization (e.g. in acorn ants, Diamond et al., 2018), these methods are time and resource consuming, and therefore can only be applied in a limited number of cities or species that are experimentally tractable. Alternatively, sequencing approaches can provide cost effective and detailed information on the molecular evolution in numerous cities simultaneously. For example, a combination of molecular based studies demonstrated independent acquisition of resistance to polychlorinated biphenyls (PCBs) involving the same physiological pathways in killifish (*Fundulus heteroclitus*) in multiple city’s estuaries (Nacci et al., 2002; Reid et al., 2016; Whitehead et al., 2011).

While we often observe repeated phenotypic shifts in urban areas, they do not necessarily arise from parallel independent adaptive evolution. For example, a common garden experiment using the Virginia pepperweed (*Lepidium virginicum*) showed genetically based convergence in the urban phenotype of earlier bolting, larger size, producing fewer leaves and more seeds However, population genetic analyses revealed that urban populations were likely derived from the same inbred haplotype, demonstrating that such parallelism in urban phenotype was facilitated by extensive gene flow among cities combined with lineage sorting within habitats, as opposed to multiple independent evolutionary events (Yakub & Tiffin, 2017). Similarly, in the Gulf killifish (*Fundulus grandis*), adaptive resistance to industrial pollutants was found in four populations, and was shown to be the result of a *de novo* mutation in only one of the populations, the three other population likely benefiting from a variant selected from their standing genetic variation or introgression (Lee & Coop, 2017). In both cases, the adaptive parallelism was facilitated by demographic events, highlighting the need for combined analyses that explore both adaptive and non-adaptive evolution to investigate how urbanization shapes evolutionary trajectories. The parallelism of non-adaptive processes has been particularly ignored in studies, even though theory suggests that genetic drift and gene flow can lead to parallel evolutionary patterns (e.g. Santangelo et al., 2018). Our results suggest that contrary to the prevailing hypothesis, cities can facilitate the demographic spread, growth and adaptation of species, especially for those with a cosmopolitan distribution.

## Acknowledgements

We would like to thank Sophie Koch, Inder Sheoran, and Beata Cohan for performing DNA extractions and making the genomic libraries, and all the GLUE collaborators for collecting clover around the world. This research was funded by CRC, School of Cities grant to AEC, NSERC Discovery grants to RWN and MTJJ and NSERC EWR Steacie Award to MTJJ.

## Data Accessibility Statement

All code is available on AEC’s Github page. BAM files have been deposited in the European Nucleotide Archive (ENA BioProject PRJEB48967).

## Notes

### Competing Interest Statement

The authors have declared no competing interest.

## REFERENCES

1. Beninde, J., Feldmeier, S., Veith, M., & Hochkirch, A. (2018). Admixture of hybrid swarms of native and introduced lizards in cities is determined by the cityscape structure and invasion history. Proceedings of the Royal Society B, 285(1883), 20180143. https://doi.org/10.1098/rspb.2018.0143

2. Björklund, M., Ruiz, I., & Senar, J. C. (2009). Genetic differentiation in the urban habitat: The great tits (*Parus major*) of the parks of Barcelona city. Biological Journal of the Linnean Society, 99(1), 9–19. https://doi.org/10.1111/j.1095-8312.2009.01335.x

3. Caizergues, A. E., Le Luyer, J., Grégoire, A., Szulkin, M., Senar, J.-C., Charmantier, A., & Perrier, C. (2022). Epigenetics and the city: Non-parallel DNA methylation modifications across pairs of urban-forest Great tit populations. Evolutionary Applications, 15(1), 149–165. https://doi.org/10. 1

4. Carlen, E., & Munshi-South, J. (2021). Widespread genetic connectivity of feral pigeons across the Northeastern megacity. Evolutionary Applications, 14(1), 150–162. https://doi.org/10.1111/EVA.12972

5. Combs, M., Puckett, E. E., Richardson, J., Mims, D., & Munshi-South, J. (2018). Spatial population genomics of the brown rat (*Rattus norvegicus*) in New York City. Molecular Ecology, 27(1), 83–98. https://doi.org/10.1111/mec.14437

6. Diamond, S. E., Chick, L. D., Perez, A., Strickler, S. A., & Martin, R. A. (2018). Evolution of thermal tolerance and its fitness consequences: Parallel and non-parallel responses to urban heat islands across three cities. Proceedings of the Royal Society B: Biological Sciences, 285(1882), 20180036. https://doi.org/10.1098/rspb.2018.0036

7. Evanno, G., Regnaut, S., & Goudet, J. (2005). Detecting the number of clusters of individuals using the software structure: A simulation study. Molecular Ecology, 14(8), 2611– 2620. https://doi.org/10.1111/j.1365-294X.2005.02553.x

8. Fox, E. A., Wright, A. E., Fumagalli, M., & Vieira, F. G. (2019). ngsLD: Evaluating linkage disequilibrium using genotype likelihoods. Bioinformatics, 35(19), 3855–3856. https://doi.org/10.1093/bioinformatics/btz200

9. Glenn, T. C., Nilsen, R. A., Kieran, T. J., Sanders, J. G., Bayona-Vásquez, N. J., Finger, J. W., Pierson, T. W., Bentley, K. E., Hoffberg, S. L., Louha, S., Garcia-De Leon, F. J., Del Rio Portilla, M. A., Reed, K. D., Anderson, J. L., Meece, J. K., Aggrey, S. E., Rekaya, R., Alabady, M., Belanger, M., … Faircloth, B. C. (2019). Adapterama I: Universal stubs and primers for 384 unique dual-indexed or 147,456 combinatorially-indexed Illumina libraries (iTru & iNext). PeerJ, 7, e7755. https://doi.org/10.7717/peerj.7755

10. Griffiths, A. G., Moraga, R., Tausen, M., Gupta, V., Bilton, T. P., Campbell, M. A., Ashby, R., Nagy, I., Khan, A., Larking, A., Anderson, C., Franzmayr, B., Hancock, K., Scott, A., Ellison, N. W., Cox, M. P., Asp, T., Mailund, T., Schierup, M. H., & Andersen, S. U. (2019). Breaking free: The genomics of allopolyploidy-facilitated niche expansion in white clover. The Plant Cell, 31(7), 1466–1487. https://doi.org/10.1105/TPC.18.00606

11. Grimm, N. B., Faeth, S. H., Golubiewski, N. E., Redman, C. L., Wu, J., Bai, X., & Briggs, J. M. (2008). Global ghange and the ecology of cities. Science, 319(5864), 756–760. https://doi.org/10.1126/science.1150195

12. Gutenkunst, R., Hernandez, R., Williamson, S., & Bustamante, C. (2010). Diffusion approximations for demographic inference: DaDi. Nature Precedings, 1–1. https://doi.org/10.1038/npre.2010.4594.1

13. Han, E., Sinsheimer, J. S., & Novembre, J. (2014). Characterizing bias in population genetic inferences from low-coverage sequencing data. Molecular Biology and Evolution, 31(3), 723–735. https://doi.org/10.1093/MOLBEV/MST229

14. Hanghøj, K., Moltke, I., Andersen, P. A., Manica, A., & Korneliussen, T. S. (2019). Fast and accurate relatedness estimation from high-throughput sequencing data in the presence of inbreeding. GigaScience, 8(5), giz034. https://doi.org/10.1093/gigascience/giz034

15. Hanski, I. (1998). Metapopulation dynamics. Nature, 396(6706), 41–49. https://doi.org/10.1038/23876

16. Hedrick, P. W., & Lacy, R. C. (2015). Measuring relatedness between inbred individuals. Journal of Heredity, 106(1), 20–25. https://doi.org/10.1093/jhered/esu072

17. Hudson, R. R., Slatkin, M., & Maddison, W. P. (1992). Estimation of levels of gene flow from DNA sequence data. Genetics, 132(2), 583–589. https://doi.org/10.1093/genetics/132.2.583

18. Johnson, M. T. J., & Munshi-South, J. (2017). Evolution of life in urban environments. Science, 358(6363), eaam8327. https://doi.org/10.1126/science.aam8327

19. Johnson, M. T. J., Prashad, C. M., Lavoignat, M., & Saini, H. S. (2018). Contrasting the effects of natural selection, genetic drift and gene flow on urban evolution in white clover (*Trifolium repens*). Proceedings of the Royal Society B: Biological Sciences, 285(1883), 20181019. https://doi.org/10.1098/rspb.2018.1019

20. Kakes, P. (1997). Difference between the male and female components of fitness associated with the gene *Ac* in *Trifolium repens*. Acta Botanica Neerlandica, 46(2), 219–223. https://doi.org/10.1111/plb.1997.46.2.219

21. Kjærgaard, T. (2003). A plant that changed the world: The rise and fall of clover 1000-2000. Landscape Research, 28(1), 41–49. https://doi.org/10.1080/01426390306531

22. Kopelman, N. M., Mayzel, J., Jakobsson, M., Rosenberg, N. A., & Mayrose, I. (2015). Clumpak: A program for identifying clustering modes and packaging population structure inferences across K. Molecular Ecology Resources, 15(5), 1179–1191. https://doi.org/10.1111/1755-0998.12387

23. Korneliussen, T. S., Albrechtsen, A., & Nielsen, R. (2014). ANGSD: Analysis of next generation sequencing data. BMC Bioinformatics, 15(1), 356. https://doi.org/10.1186/s12859-014-0356-4

24. Lambert, M. R., Brans, K. I., Des Roches, S., Donihue, C. M., & Diamond, S. E. (2021). Adaptive evolution in cities: Progress and misconceptions. Trends in Ecology and Evolution, 36(3), 239–257. https://doi.org/10.1016/j.tree.2020.11.002

25. Lee, K. M., & Coop, G. (2017). Distinguishing among modes of convergent adaptation using population genomic data. Genetics, 207(4), 1591–1619. https://doi.org/10.1534/genetics.117.300417 https://doi.org/10.1371/journal.pone.0037558

26. Liu, X., Huang, Y., Xu, X., Li, X., Li, X., Ciais, P., Lin, P., Gong, K., Ziegler, A. D., Chen, A., Gong, P., Chen, J., Hu, G., Chen, Y., Wang, S., Wu, Q., Huang, K., Estes, L., & Zeng, Z. (2020). High-spatiotemporal-resolution mapping of global urban change from 1985 to 2015. Nature Sustainability, 3(7), Article 7. https://doi.org/10.1038/s41893-020-0521-x

27. Lourenço, A., Álvarez, D., Wang, I. J., & Velo-Antón, G. (2017). Trapped within the city: Integrating demography, time since isolation and population-specific traits to assess the genetic effects of urbanization. Molecular Ecology, 26(6), 1498–1514. https://doi.org/10.1111/mec.14019

28. Lynch, M., Haubold, B., Pfaffelhuber, P., & Maruki, T. (2020). Inference of historical population-size changes with allele-frequency data. G3 Genes|Genomes|Genetics, 10(1), 211–223. https://doi.org/10.1534/g3.119.400854

29. McKinney, M. L. (2006). Urbanization as a major cause of biotic homogenization. Biological Conservation, 127(3), 247–260. https://doi.org/10.1016/j.biocon.2005.09.005

30. Medina, I., Cooke, G. M., & Ord, T. J. (2018). Walk, swim or fly? Locomotor mode predicts genetic differentiation in vertebrates. Ecology Letters, 21(5), 638–645. https://doi.org/10.1111/ele.12930

31. Meisner, J., & Albrechtsen, A. (2018). Inferring population structure and admixture proportions in low-depth NGS data. Genetics, 210(2), 719–731. https://doi.org/10.1534/genetics.118.301336

32. Miles, L. S., Dyer, R. J., & Verrelli, B. C. (2018). Urban hubs of connectivity: Contrasting patterns of gene flow within and among cities in the western black widow spider. Proceedings of the Royal Society B, 285(1884). https://doi.org/10.1098/RSPB.2018.1224

33. Miles, L. S., Rivkin, L. R., Johnson, M. T. J., Munshi-South, J., & Verrelli, B. C. (2019). Gene flow and genetic drift in urban environments. Molecular Ecology, 28(18), 4138– 4151. https://doi.org/10.1111/mec.15221

34. Mölder, F., Jablonski, K. P., Letcher, B., Hall, M. B., Tomkins-Tinch, C. H., Sochat, V., Forster, J., Lee, S., Twardziok, S. O., Kanitz, A., Wilm, A., Holtgrewe, M., Rahmann, S., Nahnsen, S., & Köster, J. (2021). Sustainable data analysis with Snakemake. F1000Research, 10, 33. https://doi.org/10.12688/f1000research.29032.2

35. Mueller, J. C., Kuhl, H., Boerno, S., Tella, J. L., Carrete, M., & Kempenaers, B. (2018). Evolution of genomic variation in the burrowing owl in response to recent colonization of urban areas. Proceedings of the Royal Society B, 285(1878), 20180206. https://doi.org/10.1098/rspb.2018.0206

36. Munshi-South, J., Zolnik, C. P., & Harris, S. E. (2016). Population genomics of the Anthropocene: Urbanization is negatively associated with genome-wide variation in white-footed mouse populations. Evolutionary Applications, 9(4), 546–564. https://doi.org/10.1111/eva.12357

37. Nacci, D. E., Champlin, D., Coiro, L., McKinney, R., & Jayaraman, S. (2002). Predicting the occurrence of genetic adaptation to dioxinlike compounds in populations of the estuarine fish *Fundulus heteroclitus*. Environmental Toxicology and Chemistry, 21(7), 1525–1532. https://doi.org/10.1002/etc.5620210726

38. Nei, M. (1975). Molecular population genetics and evolution. Frontiers of Biology, 40, I–288.

39. Nielsen, R., Korneliussen, T., Albrechtsen, A., Li, Y., & Wang, J. (2012). SNP calling, genotype calling, and sample allele frequency estimation from new-generation sequencing data. PLOS ONE, 7(7), e37558. https://doi.org/10.1371/journal.pone.0037558

40. Noël, S., Ouellet, M., Galois, P., & Lapointe, F.-J. (2007). Impact of urban fragmentation on the genetic structure of the eastern red-backed salamander. Conservation Genetics, 8(3), 599–606. https://doi.org/10.1007/s10592-006-9202-1

41. Noskova, E., Ulyantsev, V., Koepfli, K.-P., O’Brien, S. J., & Dobrynin, P. (2020). GADMA: Genetic algorithm for inferring demographic history of multiple populations from allele frequency spectrum data. GigaScience, 9(3), giaa005. https://doi.org/10.1093/gigascience/giaa005

42. Oziolor, E. M., Reid, N. M., Yair, S., Lee, K. M., Guberman VerPloeg, S., Bruns, P. C., Shaw, J. R., Whitehead, A., & Matson, C. W. (2019). Adaptive introgression enables evolutionary rescue from extreme environmental pollution. Science, 364(6439), 455–457. https://doi.org/10.1126/science.aav4155

43. Reid, N. M., Proestou, D. A., Clark, B. W., Warren, W. C., Colbourne, J. K., Shaw, J. R., Karchner, S. I., Hahn, M. E., Nacci, D., Oleksiak, M. F., Crawford, D. L., & Whitehead, A. (2016). The genomic landscape of rapid repeated evolutionary adaptation to toxic pollution in wild fish. Science, 354(6317), 1305–1308. https://doi.org/10.1126/science.aah4993

44. Rochat, E., Manel, S., Deschamps-Cottin, M., Widmer, I., & Joost, S. (2017). Persistence of butterfly populations in fragmented habitats along urban density gradients: Motility helps. Heredity, 119(5), Article 5. https://doi.org/10.1038/hdy.2017.40

45. Salmón, P., Jacobs, A., Ahrén, D., Biard, C., Dingemanse, N. J., Dominoni, D. M., Helm, B., Lundberg, M., Senar, J. C., Sprau, P., Visser, M. E., & Isaksson, C. (2021). Repeated genomic signatures of adaptation to urbanisation in a songbird across Europe. Nature Communications, 12(1), 1–14. https://doi.org/10.1038/s41467-021-23027-w

46. Santangelo, J. S., Miles, L. S., Breitbart, S. T., Murray-Stoker, D., Rivkin, L. R., Johnson, M. T. J., & Ness, R. W. (2020). Urban environments as a framework to study parallel evolution. In M. Szulkin, J. Munshi-South, & A. Charmantier (Eds.), Urban Evolutionary Biology (pp. 36–53). Oxford University Press.

47. Santangelo, J. S., Ness, R. W., Cohan, B., Fitzpatrick, C. R., Innes, S. G., Koch, S., Miles, L. S., Munim, S., Peres-Neto, P. R., Prashad, C., Tong, A. T., Aguirre, W. E., Akinwole, P. O., Alberti, M., Álvarez, J., Anderson, J. T., Anderson, J. J., Ando, Y., Andrew, N. R., … Johnson, M. T. J. (2022). Global urban environmental change drives adaptation in white clover. Science, 375(6586), 1275–1281. https://doi.org/10.1126/science.abk0989

48. Santangelo, J. S., Rivkin, L. R., Advenard, C., & Thompson, K. A. (2020). Multivariate phenotypic divergence along an urbanization gradient. Biology Letters, 16(9), 20200511. https://doi.org/10.1098/rsbl.2020.0511

49. Santangelo, J. S., Rivkin, L. R., & Johnson, M. T. J. (2018). The evolution of city life. Proceedings of the Royal Society B: Biological Sciences, 285(1884), 20181529. https://doi.org/10.1098/rspb.2018.1529

50. Schmidt, C., Domaratzki, M., Kinnunen, R. P., Bowman, J., & Garroway, C. J. (2020). Continent-wide effects of urbanization on bird and mammal genetic diversity. Proceedings of the Royal Society B, 287(1920), 20192497. https://doi.org/10.1098/rspb.2019.2497

51. Skotte, L., Korneliussen, T. S., & Albrechtsen, A. (2013). Estimating individual admixture proportions from next generation sequencing data. Genetics, 195(3), 693–702. https://doi.org/10.1534/genetics.113.154138

52. Somers, C. M., McCarry, B. E., Malek, F., & Quinn, J. S. (2004). Reduction of particulate air pollution lowers the risk of heritable mutations in mice. Science, 304(5673), 1008– 1010. https://doi.org/10.1126/science.1095815

53. Tajima, F. (1989). The effect of change in population size on DNA polymorphism. Genetics, 123(3), 597–601. https://doi.org/10.1093/genetics/123.3.597

54. Tajima, F. (1983). Evolutionary relationship of DNA sequences in finite populations. Genetics, 105(2), 437–460. https://doi.org/10.1093/genetics/105.2.437111/eva.13334

55. Theodorou, P., Radzevičiūtė, R., Kahnt, B., Soro, A., Grosse, I., & Paxton, R. J. (2018). Genome-wide single nucleotide polymorphism scan suggests adaptation to urbanization in an important pollinator, the red-tailed bumblebee (*Bombus lapidarius L*.). Proceedings of the Royal Society B: Biological Sciences, 285(1877). https://doi.org/10.1098/rspb.2017.2806

56. Wenzel, A., Grass, I., Belavadi, V. V., & Tscharntke, T. (2020). How urbanization is driving pollinator diversity and pollination – A systematic review. Biological Conservation, 241, 108321. https://doi.org/10.1016/j.biocon.2019.108321

57. Whitehead, A., Pilcher, W., Champlin, D., & Nacci, D. (2011). Common mechanism underlies repeated evolution of extreme pollution tolerance. Proceedings of the Royal Society B: Biological Sciences, 279(1728), 427–433. https://doi.org/10.1098/rspb.2011.0847

58. Williams, W. M., Mason, K.M., & Williamson, M. L. (1998). Genetic analysis of shikimate dehydrogenase allozymes in *Trifolium repens L*. Theor Appl Genet, 96, 859–868.

59. Wright, S. (1984). Evolution and the Genetics of Populations, Volume 4: Variability Within and Among Natural Populations. University of Chicago Press.

60. Wu, F., Ma, S., Zhou, J., Han, C., Hu, R., Yang, X., Nie, G., & Zhang, X. (2021). Genetic diversity and population structure analysis in a large collection of white clover (*Trifolium repens L*.) germplasm worldwide. PeerJ, 9, e11325. https://doi.org/10.7717/peerj.11325

61. Yakub, M., & Tiffin, P. (2017). Living in the city: Urban environments shape the evolution of a native annual plant. Global Change Biology, 23(5), 2082–2089. https://doi.org/10.1111/gcb.13528

62. Yauk, C., Polyzos, A., Rowan-Carroll, A., Somers, C. M., Godschalk, R. W., Van Schooten, F. J., Berndt, M. L., Pogribny, I. P., Koturbash, I., Williams, A., Douglas, G. R., & Kovalchuk, O. (2008). Germ-line mutations, DNA damage, and global hypermethylation in mice exposed to particulate air pollution in an urban/industrial location. Proceedings of the National Academy of Sciences, 105(2), 605–610. https://doi.org/10.1073/pnas.0705896105

