## Supplementary material for "Does urbanization lead to parallel demographic shifts across the world in a cosmopolitan plant?"

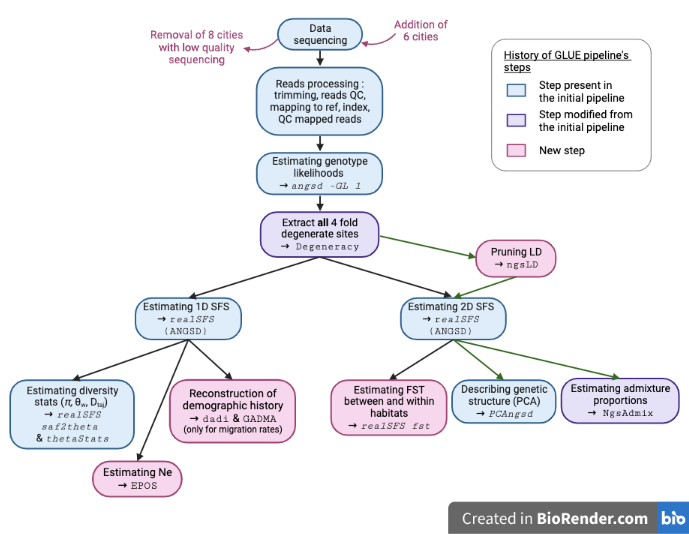

**Figure S1:** Flowchart of the bioinformatic pipeline’s steps used for the analysis presented in the current paper. Each step is represented in a color indicating if it was reused (blue) or modified (purple) from the Santangelo’s et al (2021) pipeline, or added (pink) in the current pipeline. Note that even if steps are colored in blue (meaning the scripts were reused from Santangelo et al 2021), since the dataset was updated and some upstream steps were refined, the outcome of the analyses are always quantitatively different from the previously published work. Green arrows represent an alternative path of analyses that included a filtration of linked positions. This is a simplified flowchart including only the main steps of the pipeline, for detailed steps and functions see the publicly available and reproducible pipeline on AEC’s github.

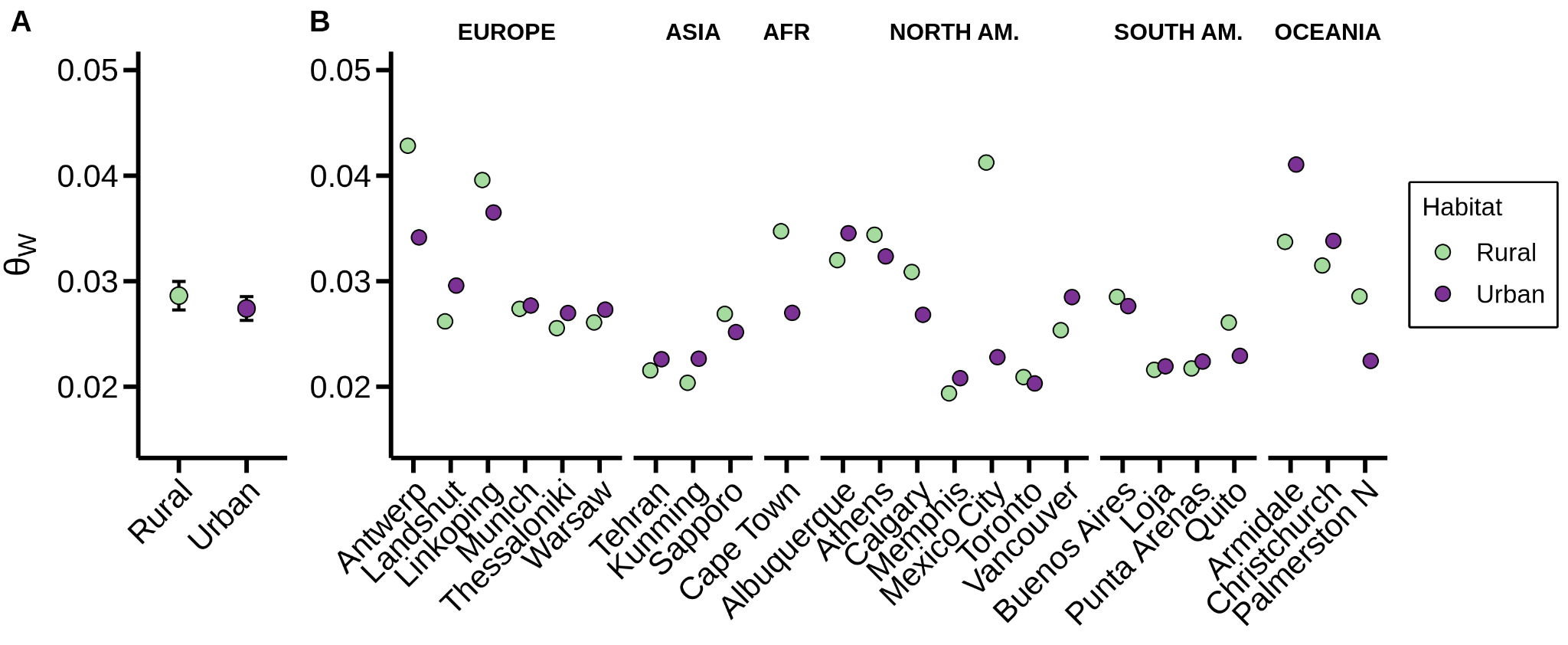

**Figure S2:** Genome wide nucleotide diversity measure with Watterson’s θ. A) Rural and urban θ_w_ averaged on the overall dataset (+-SE). B) Per city per habitat θ_w_ (note that 95% CI intervals are plotted but small enough to be hidden behind each dot).

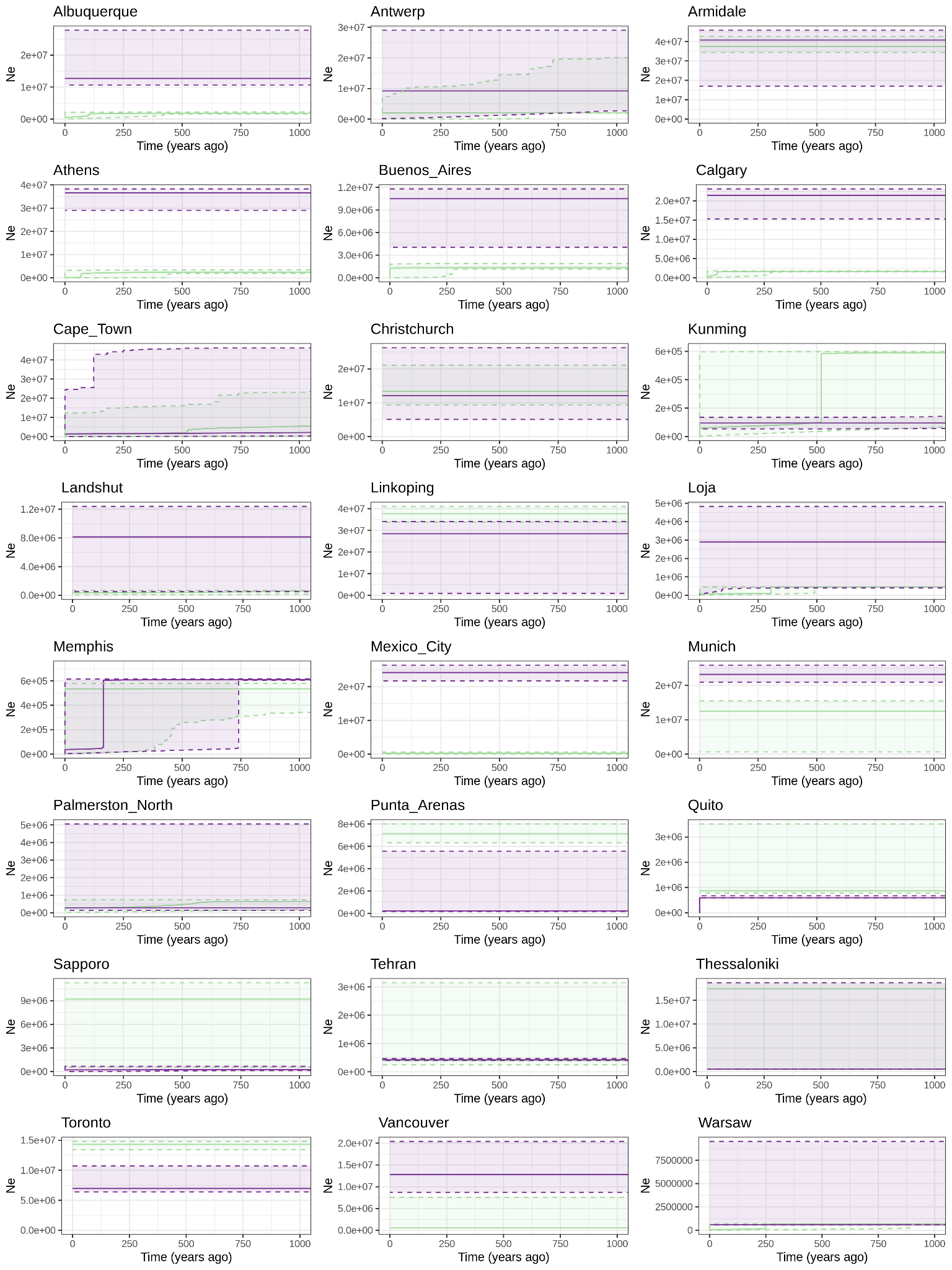

**Figure S3:** Effective population size reconstruction with EPOS. Rural (green) and urban (purple) solid lines represent the estimated median and the dashed lines represent 5% and 95% quantiles.

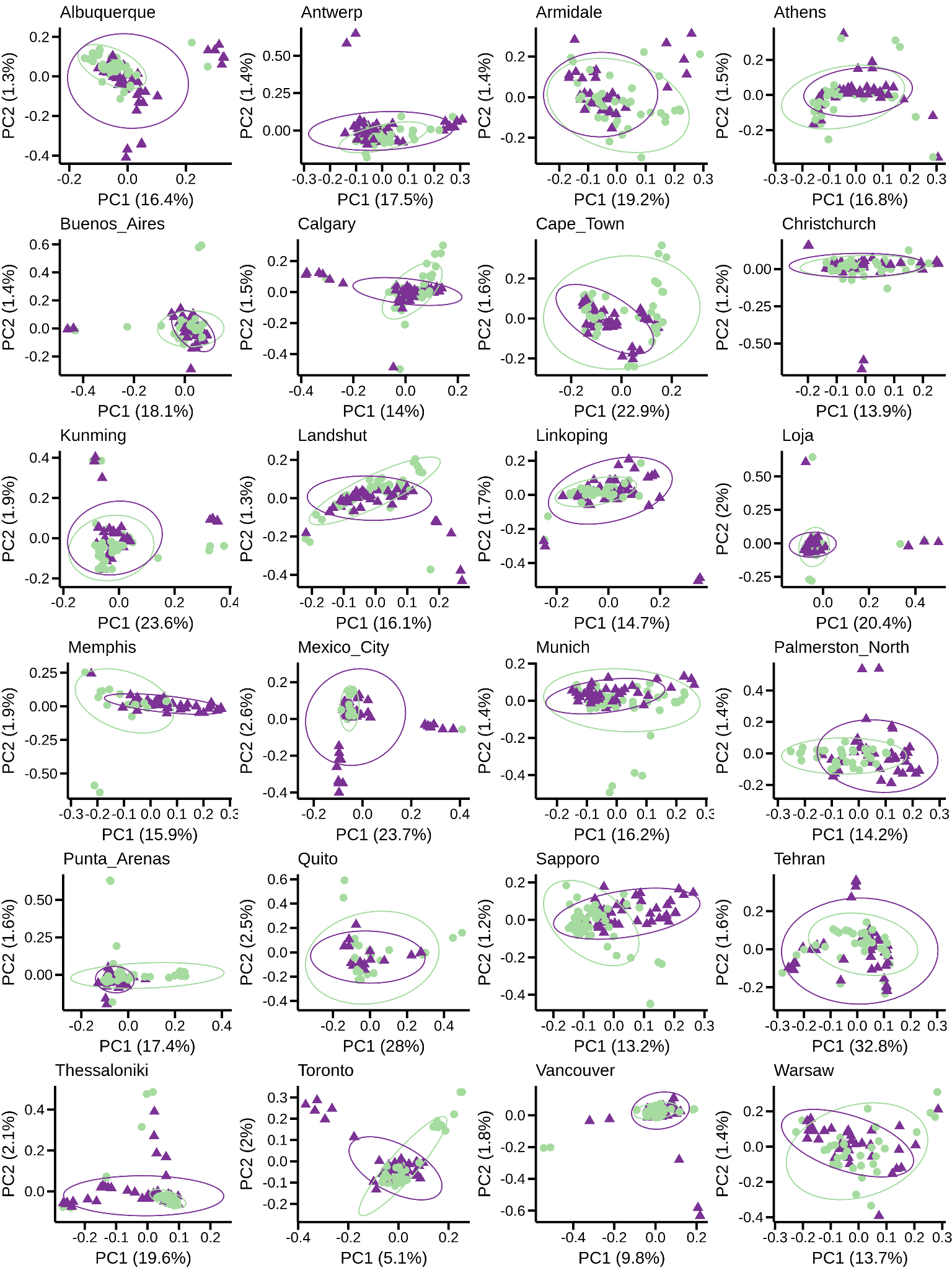

**Figure S4:** Genetic structure between urban and rural individuals revealed by per city principal component analyses (PCA). Overall, we observe little genetic clustering between urban (purple triangles) and rural (green dots), as demonstrated by the large overlap of 95% CI ellipses.

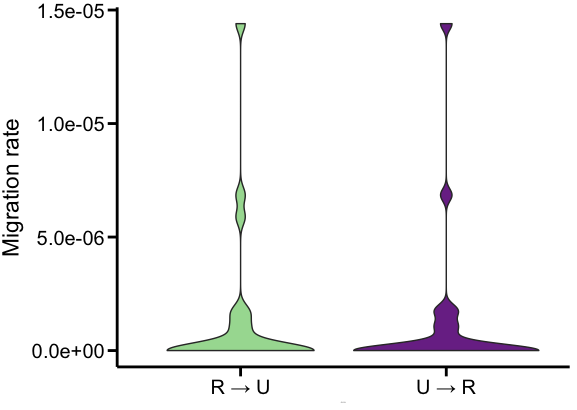

**Figure S5:** Migration rates in the Rural → Urban and Urban → Rural directions estimated with the demographic history reconstruction via *dadi*. The unit of migration is defined as the proportion of chromosomes that are new migrants in a population per generation.

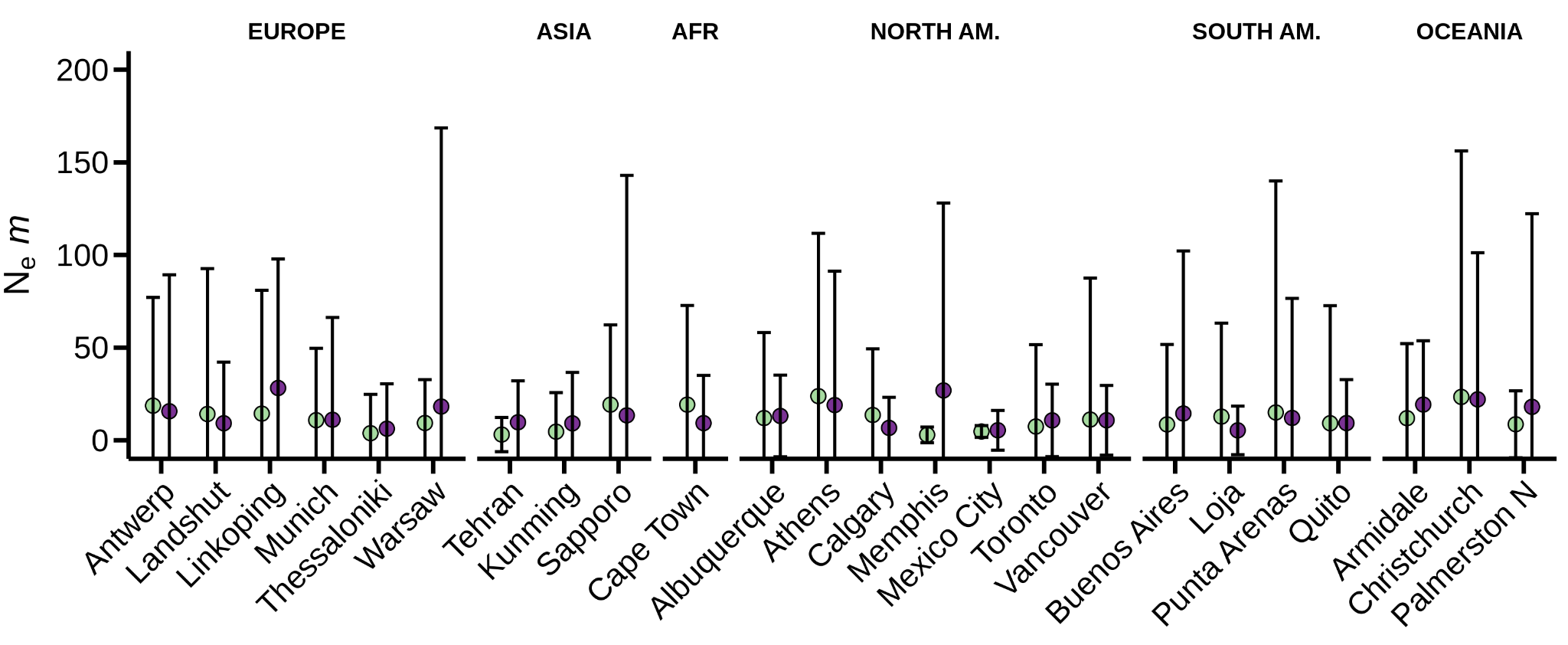

**Figure S6:** N_e_*m* (Effective population size x migration rate) computed from F_ST_ values following Wright (1931) as F_ST_= 1/(4*N_e_m*+1).
